# Serotonin shapes the temporal window for associative fear learning

**DOI:** 10.1101/2025.08.06.669013

**Authors:** Yulin Zhao, Xuelin Li, Tianyi Chang, Shu Xie, Yiling Wu, Shaochuang Li, Jiayi Cao, Di Wang, Fei Deng, Jiesi Feng, Yanyi Huang, Yulong Li

## Abstract

Fear learning is a critical adaptive mechanism that enables the association of an environmental cue (the conditioned stimulus, CS) with a potential threat (the unconditioned stimulus, US), even when these events are temporally discontiguous. When associations are instead formed between unrelated cues and threats, the maladaptive learning process can give rise to neurological disorders. Therefore, maintaining accurate temporal boundaries for forming such associations is crucial for distinguishing real threats from irrelevant stimuli. Despite its importance, the mechanisms defining this precise time window remain poorly understood. Here, we report that serotonin plays a crucial role in modulating the associable interval in trace fear conditioning. Specifically, we show that serotonergic projections from the dorsal raphe nuclei (DRN) to the CA1 region of the ventral hippocampus (vCA1) dynamically regulate the association of discontiguous CS and US. Mechanistically, we demonstrate that sustained serotonin release in the vCA1 maintains the ensemble activity of 5-HT2C receptor-expressing pyramidal neurons, facilitating the association between temporally separated events. In contrast, when the trace interval exceeds the associable window, transient serotonin release fails to drive vCA1 neuronal ensemble activity, thus preventing maladaptive learning. These findings highlight a key role for serotonin in regulating the temporal precision of fear learning and reveal the DRN-vCA1 serotonergic pathway as a potential therapeutic target for related disorders.

## Main

In animals, associative fear learning is essential for survival. To better adapt to the complex environments, animals have evolved the ability to form associations between an environmental cue (the conditional stimulus, CS) and a dangerous situation (the unconditional stimulus, US) that are not temporally synchronous. This phenomenon is often modeled in the laboratory as Pavlovian trace fear conditioning, a paradigm widely used to study adaptive fear learning^1–5^. Developing an accurate association within an appropriate time window (i.e., trace interval) is crucial to avoid maladaptive learning, which can lead to anxiety-related disorders such as post-traumatic stress disorder^6^. Previous studies characterized the brain regions involved in trace fear conditioning. Lesions of the hippocampus have been shown to attenuate fear responses in trace fear conditioning^7–9^. In addition, the prefrontal cortex (PFC) has been implicated in the association of temporally separated events^10,11^. As a key brain region in fear learning, pretraining lesions of the basolateral amygdala have also been reported to reduce fear expression in trace fear conditioning^12^. Despite these findings, the specific molecular and circuit mechanisms that regulate the associable interval remain unclear.

The serotonergic system is implicated in myriad brain functions in mammals, including cognition and emotional processing^13^. In humans, mutations in serotonin receptors (5- HTRs) or declined serotonin transporter availability are associated with learning impairments and an increased risk for affective disorders^14,15^. Clinical evidence also indicates that selective serotonin reuptake inhibitors (SSRIs) may produce early adverse effects in certain individuals^16^. In a study involving healthy volunteers, depletion of tryptophan (the precursor in serotonin synthesis) revealed the importance of serotonin in memory formation^17^. Moreover, rodent studies suggest that serotonin is a critical neuromodulator of pathological fear learning^18–20^; however, the role of serotonin in trace fear conditioning remains poorly understood. Notably, studies have shown that serotonergic neurons project widely to the forebrain, including regions implicated in trace fear conditioning, such as the hippocampus, the basal amygdala (BA), and the PFC^21–23^.

In this study, we show that manipulating serotonin levels modulates the associable trace interval. Using a genetically encoded serotonin sensor, we identified specific patterns of serotonergic dynamics in the CA1 region of the ventral hippocampus (vCA1) during associable trace fear conditioning. Furthermore, we found that this regulation is mediated by serotonergic projections from the dorsal raphe nuclei (DRN) to the vCA1. Using *in vivo* calcium recording combined with activity-dependent labeling, we demonstrated that sustained serotonin release in the vCA1 during associable trace conditioning acts via 5-HT2C receptors to modulate the activity of pyramidal neuronal ensembles; in contrast, transient serotonin release during non-associable trace conditioning fails to activate vCA1 neuronal ensembles, thereby preventing the formation of a CS-US association.

## Results

### Serotonin regulates the associable window in trace fear conditioning

To characterize the associable window in mice trace fear conditioning, we utilized the classical Pavlovian cued fear conditioning model^24,25^ and systematically increased the trace interval between the CS and US; freezing behavior in response to the CS during the test session served as an indicator of a successful CS-US association (Fig. 1a). To minimize confounding effects from repeated training and/or the intertrial interval, each mouse in our study underwent only one training trial. On the test day, we observed that the freezing behavior decreased as the trace interval increased (Fig. 1b). Notably, mice exhibited no apparent freezing behavior (indicated by the absence of a significant difference compared to baseline) when the trace interval reached 60 seconds (Fig.1b).

**Fig. 1.**
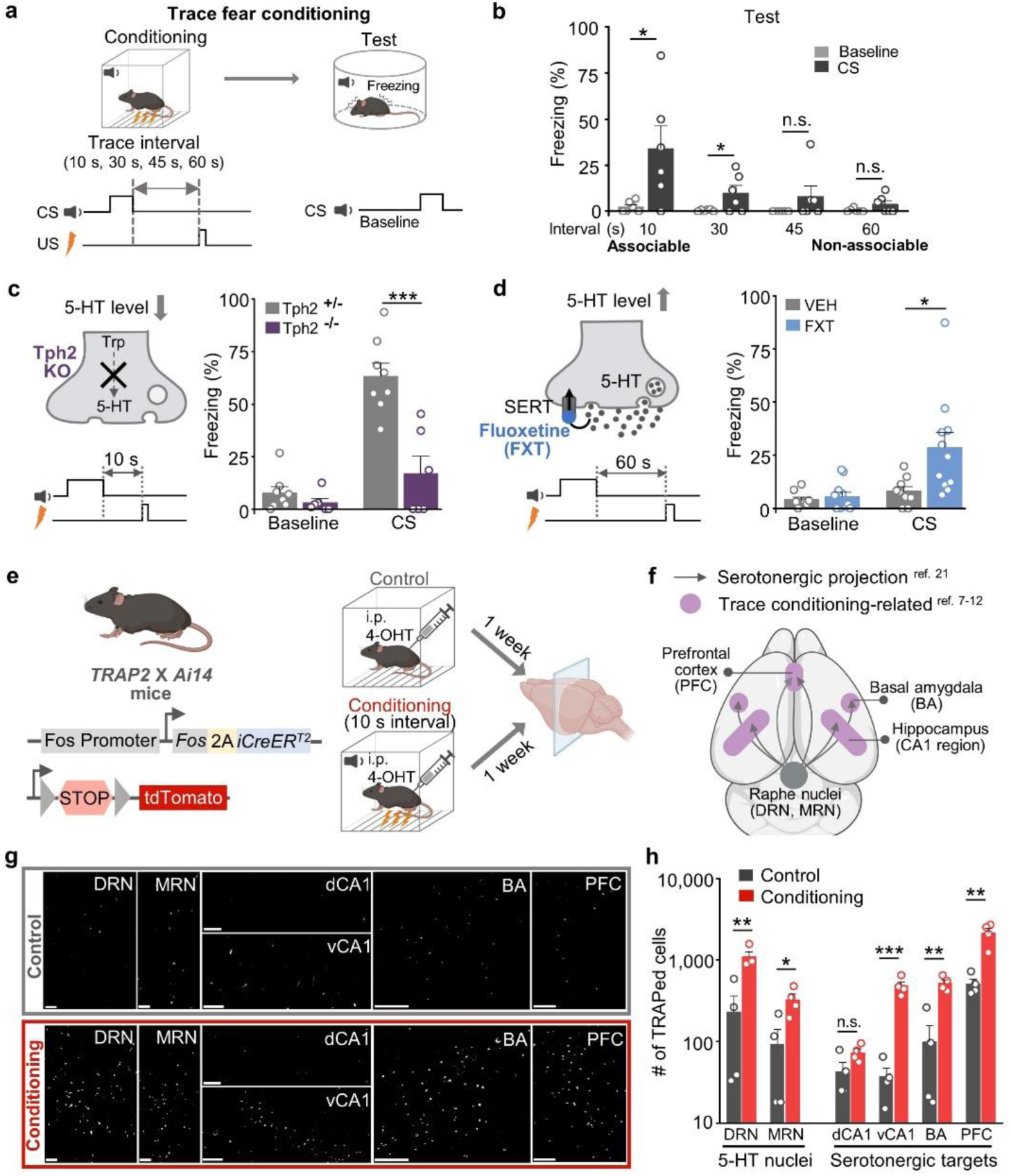
Serotonin regulates the associable window in trace fear conditioning. a, Schematic of the trace fear conditioning paradigm: mice underwent single-trial conditioning with increasing trace intervals. Freezing behavior in response to the CS was assessed during the test. **b,** Summary of freezing time (%) measured during baseline or CS presentation with indicated trace intervals. Baseline vs. CS, * *p*=0.0377 (10 s), * *p*=0.0455 (30 s), n.s. (45 s, 60 s); paired *t* test; n=6 mice/group. **c,** Schematic diagram (left) and summary (right) of freezing time (%) of both homozygous and heterozygous Tph2 KO mice trained with 10-s interval conditioning. *** *p*=0.0007; unpaired *t* test; n=6-8 mice/group. **d,** Schematic diagram (left) and summary (right) of freezing time (%) in mice treated with 10 mg/kg fluoxetine (FXT) or vehicle (VEH) and trained with 60-s interval conditioning. * *p*=0.0212; unpaired *t* test; n=9-11 mice/group. **e,** *TRAP2 x Ai14* mice were used for activity-dependent neuronal labeling (tdTomato). Mice received 50 mg/kg 4-OH tamoxifen (4-OHT) before training. Conditioning mice were trained with 10-s interval conditioning; controls were exposed to the chamber without CS/US. **f,** Schematic of brain regions involved in trace fear conditioning and serotonergic projections from the raphe nuclei. **g,** Example images of TRAPed cells (white dots) in the indicated brain regions; scale bars, 200 μm. **h,** Summary of TRAPed cells counts in control (black, n=4) and conditioning (red, n=4) groups. Regions: DRN (** *p*=0.0059), MRN (* *p*=0.0239), dCA1 (n.s.), vCA1 (*** *p*=0.0004), BA (** *p*=0.0016), and PFC (** *p*=0.0029); unpaired t-test. Data shown as means ± SEM.

Next, to examine the role of serotonin in the associable window of trace fear conditioning, we manipulated serotonin levels either by knocking out serotonin synthesis enzyme, tryptophan hydroxylase 2 (Tph2), or by applying the SSRI, fluoxetine^26^. We found that with a 10-s trace interval, homozygous Tph2 knockout (KO) mice exhibited significantly reduced freezing behavior compared to heterozygous KO mice (Fig. 1c). In contrast, fluoxetine-treated wild-type mice displayed significantly increased freezing behavior compared to vehicle-treated controls with 60-s trace interval, a condition in which mice typically fail to form CS-US associations (Fig. 1d). Interestingly, fluoxetine treatment had no effect on learning efficiency with a 10-s trace interval (Extended Data Fig. 1a & 1b). Moreover, fluoxetine treatment right after the training phase had no effect on freezing behavior, indicating that the effect of fluoxetine was not attributed to its effect on the memory consolidation (Extended Data Fig. 1c & 1d). To rule out the possibility that fluoxetine-induced increase in freezing behavior was due to elevated anxiety, we conducted elevated plus maze and open-field tests and found that fluoxetine-treated mice did not exhibit anxiety-like behaviors in these tests (Extended Data Fig. 1e-h). Notably, similar results regarding associable window and serotonin-dependent regulation measured in male mice were also observed in females, indicating no sexual dimorphism in this behavior (Extended Data Fig. 2a-d). Taken together, these results indicate that serotonin plays an essential role in regulating the associable window in rodent trace fear conditioning.

### US-induced sustained serotonin release in the vCA1 correlates with the associability of trace fear conditioning

To dissect the specific serotonergic projections that regulate trace fear conditioning, we performed activity-dependent labeling experiments and analyzed the brain regions that are both innervated by serotonergic neurons and are known to be involved in the CS-US association during trace fear conditioning. These regions include the PFC, the dorsal CA1 region of the hippocampus (dCA1), the vCA1, and the BA^7–12^. We used the targeted recombination in active population 2 (*TRAP2*) x *Ai14* mouse to label neurons that are active during training with tdTomato in the presence of 4-hydroxytamoxifen (4-OHT)^27^ (Fig. 1e). Mice that underwent fear conditioning with a 10-s interval showed significantly more TRAPed (i.e., labeled) cells in all brain regions analyzed, with the sole exception of dCA1, compared to control group (Fig. 1f-h).

Next, to determine whether the brain regions identified above receive serotonergic regulation during trace fear conditioning, we took advantage of our recently developed serotonin sensor, 5-HT3.0^(ref.28)^. We injected adeno-associated viruses (AAVs) expressing the 5-HT3.0 sensor into the vCA1, BA, and PFC, and then recorded serotonin dynamics with fiber photometry with training intervals of either 10-s or 60-s (Fig. 2a). All three brain regions exhibited shock-induced serotonin signals, whereas no detectable signal was induced by the tone regardless of the trace interval. Additionally, we found no significant difference in the peak serotonin signals during US presentation between the 10-s and 60-s intervals (Fig. 2b-e). Interestingly, we observed a prolonged serotonin signal in the vCA1 in response to the US during the 10-s trace interval and this signal persisted well after stimuli cessation. In contrast, the serotonin signal in the vCA1 induced with a 60-s interval was transient, resembling the patterns observed in other brain regions (Fig. 2c & 2d). Quantitative analysis revealed that the normalized area under the curve (AUC) for the US-induced serotonin signals in the vCA1 during 10-s conditioning was significantly higher compared to other brain regions and 60-s conditioning (Fig. 2f-h). Interestingly, no CS-evoked serotonin release was measurable during the test (Extended Data Fig. 3a-c). To evaluate whether the sustained serotonin release is specific to associable trace fear conditioning, we recorded serotonin dynamics in the vCA1 and BA during delay fear conditioning, in which the CS and the US overlap and terminate concurrently^3^ (Extended Data Fig. 3d). Both regions showed transient serotonin signals in response to the US during delay fear conditioning (Extended Data Fig. 3e-g).

**Fig. 2.**
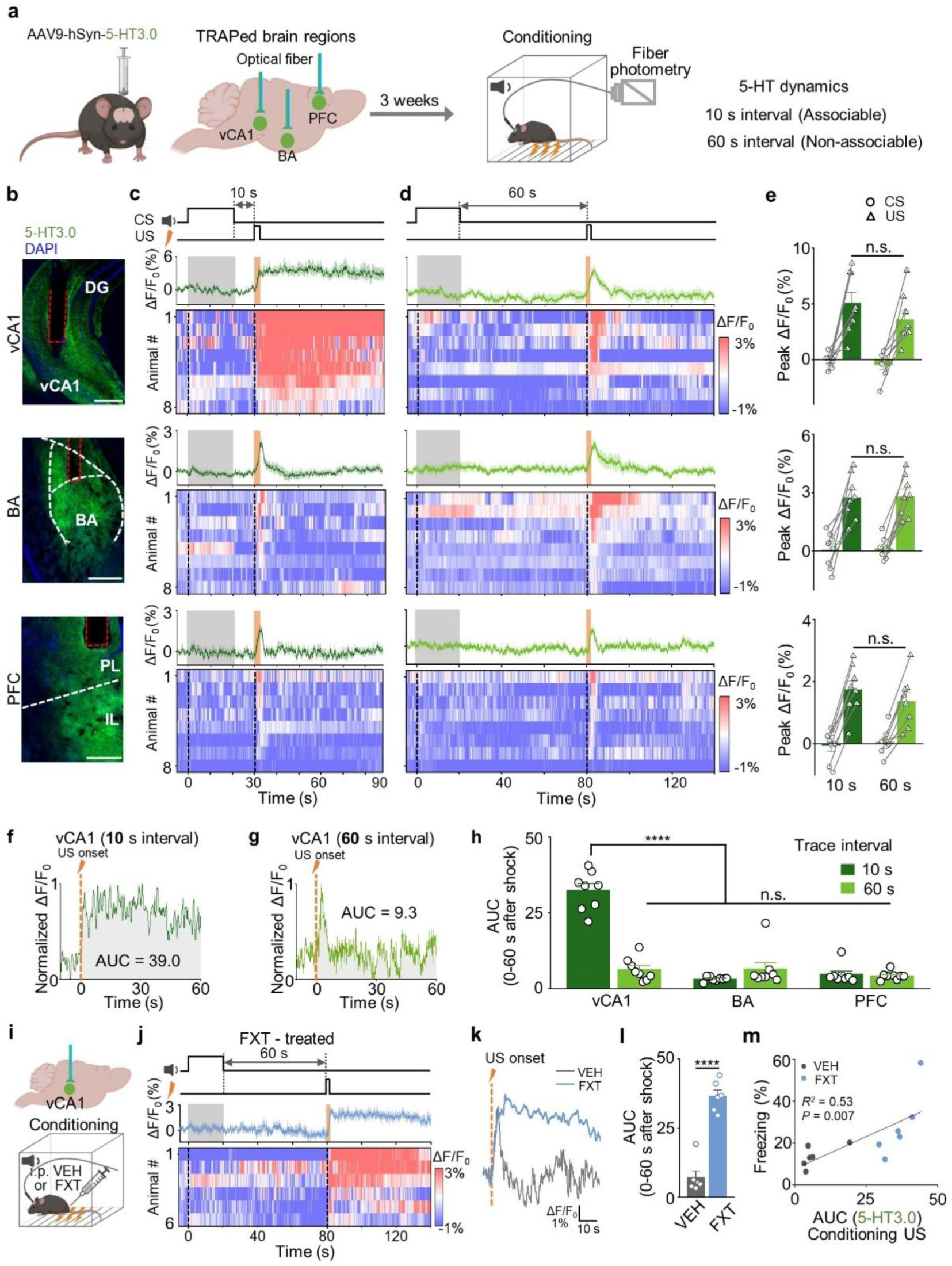
Sustained serotonin release in the vCA1 correlates with the associability of trace fear conditioning. a, AAV9-mediated expression of the serotonin sensor 5- HT3.0 in the vCA1, BA, and PFC. Optical fibers were implanted for fiber photometry recordings during associable (10-s) or non-associable (60-s) interval training. **b,** 5- HT3.0 expression in the vCA1, BA, and PFC. Red dotted lines indicate fiber tracks. White dotted lines show boundary of indicated brain region. Scale bars, 500 μm. **c & d,** 5-HT3.0 signals measured in the vCA1, BA, and PFC during 10-s (**c**) or 60-s (**d**) conditioning. Upper traces: averaged 5-HT3.0 signals; the grey and orange shaded areas indicate the CS and US delivery, respectively. Lower heatmaps: the 5-HT3.0 signals for individual mice; the dashed vertical lines mark the stimuli onset. **e,** Summary of peak ΔF/F0 during CS (circles, 20 s) or US (triangles, 2 s) delivery. Peak ΔF/F0 was calculated by averaging 10 data points near the peak. **f & g,** Example traces of normalized ΔF/F0 60 s after US delivery in 10-s interval (**f**) or 60-s interval (**g**) conditioning; the dashed lines indicate US delivery. Shaded regions denote the area under curve (AUC) analysis. **h,** Summary of AUC for the 5-HT3.0 signals in the vCA1, BA, and PFC during 10-s or 60-s interval conditioning. **** *p*<0.0001; One-way ANOVA with Tukey’s post hoc test; n=8 mice/group. **i,** 5-HT3.0 expression and fiber implantation in vCA1 for recording serotonin dynamics after 10 mg/kg FXT or VEH treatment. **j,** Averaged trace (top) and heatmap (bottom) of 5-HT3.0 signals during 60 s interval conditioning. **k,** Example 5-HT3.0 traces upon US delivery in VEH (black) and FXT (blue) groups. **l,** Summary of AUC for the 5-HT3.0 signals in the indicated groups. **** *p*<0.0001, unpaired *t test;* n=4 mice/group. **m,** Correlation between freezing behavior and the AUC for the 5-HT3.0 signals after US delivery; the Pearson’s correlation coefficient is shown. Data shown as means ± SEM.

Given that fluoxetine treatment enhanced the CS-US association during 60-s trace conditioning, we then tested whether this effect was accompanied by changes in serotonin signaling in the vCA1 (Fig. 2i). We found that fluoxetine treatment induced a sustained US-induced serotonin signal in the vCA1 (Fig. 2j & 2k), in line with increased freezing behavior (Fig. 2l & 2m). Since we observed similar behavioral results between male and female mice in response to manipulating serotonin levels, we also recorded serotonin signals in females. Similar to male mice, the serotonin signals in the vCA1 of female mice exhibited a sustained increase to the US only during 10-s trace fear conditioning. Moreover, fluoxetine treatment also induced a sustained serotonin signal under 60-s trace conditioning (Extended Data Fig. 4a-e).

### Serotonergic projections from the DRN to the vCA1 are critical for establishing the CS-US association in trace fear conditioning

To directly determine whether serotonin release in the vCA1 is essential for establishing the CS-US association in trace fear conditioning, we employed optogenetic tools in *Sert-Cre* mice^29^ to manipulate the activity of serotonergic projections to the vCA1 with high temporal resolution (Fig. 3a & Extended Data Fig. 5c). We initially tested the learning ability of *Sert-Cre* mice and confirmed that their freezing levels during 10-s trace conditioning were comparable to those of wild-type mice (Extended Data Fig. 5a & 5b). While previous tracing studies in rodents suggest that the hippocampus predominantly receives serotonergic projections from the median raphe nuclei (MRN)^30^, recent functional studies showed that the DRN also innervates the hippocampus, including the ventral hippocampus^31–33^. To test whether serotonergic inputs to the vCA1 are required for CS-US association, we injected Cre-dependent AAVs expressing eNpHR3.0 (a yellow light-sensitive inhibitory opsin)^34^ into the DRN or the MRN of *Sert-Cre* mice and delivered 556 nm constant light into the vCA1 during 10 s trace conditioning to inhibit serotonergic terminals (Fig. 3b & Extended Data Fig. 5d). Photoinhibition of DRN-vCA1 serotonergic projections throughout the training process significantly reduced freezing behavior in eNpHR3.0-expressing mice compared to mCherry-expressing controls (Fig. 3c). In contrast, photoinhibition of MRN-vCA1 serotonergic projections has no effect (Extended Data Fig. 5e).

**Fig. 3.**
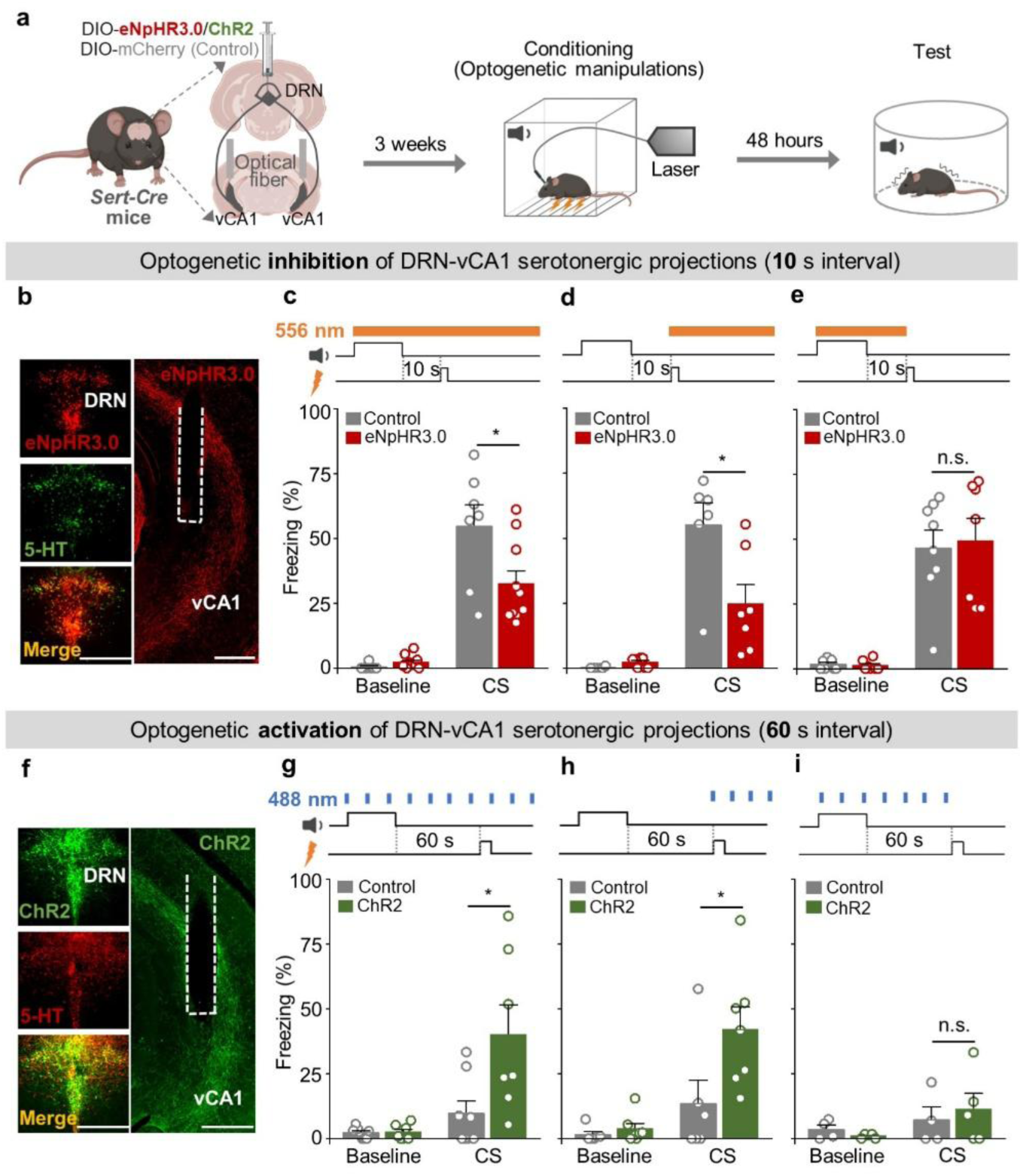
The DRN-vCA1 serotonergic circuit modulates the associability of trace fear conditioning. **a.** Left: schematic diagram of AAV-DIO-eNpHR3.0, AAV-DIO-ChR2, and AAV-DIO-mCherry injection into the DRN of *Sert-Cre* mice, with optical fiber implantation in the bilateral vCA1. Right: the experimental design of optogenetic manipulation during trace fear conditioning. **b & f.** Example images of eNpHR3.0 (**b**) or ChR2 (**f**) colocalization with 5-HT in the DRN (left), and eNpHR3.0 (**b**) or ChR2 (**f**) expression in the vCA1 (right). Scale bars, 500 μm. **c & g.** Summary of freezing time after optogenetic inhibition (**c**) or activation (**g**) of the DRN-vCA1 serotonergic circuit during the entire conditioning session. * *p*=0.0299 (eNpHR3.0 vs. control), * *p*=0.0252 (ChR2 vs. control); unpaired *t* test; n=7-10 mice/group. **d & h.** Summary of freezing time after optogenetic inhibition (**d**) or activation (**h**) of the DRN-vCA1 serotonergic circuit upon shock delivery. * *p*=0.0203 (eNpHR3.0 vs. control), * *p*=0.0467 (ChR2 vs. control); unpaired *t* test; n=6-7 mice/group. **e & i.** Summary of freezing time after optogenetic inhibition (**e**) or activation (**i**) of the DRN-vCA1 serotonergic circuit during the tone and the interval. n.s.; unpaired *t* test; n=4-8 mice/group. Data shown as means ± SEM.

To specifically assess whether the US-induced sustained serotonin release in the vCA1 is required for establishing the CS-US association, we delivered 556 nm light simultaneously with the presence of the US. This photoinhibition significantly reduced freezing behavior in eNpHR3.0-expressing mice (Fig. 3d), whereas photoinhibition applied before the US had no effect (Fig. 3e). In parallel optogenetic activation experiments, mice expressing ChR2 (a blue light sensitive excitatory opsin)^35^ in DRN- vCA1 serotonergic terminals displayed significantly increased freezing behavior compared to control mice when the activating light pulses were delivered either throughout training or specifically during the US, but not when applied before the US (Fig. 3f-i). Similarly, chemogenetic activation of DRN-vCA1 serotonergic neurons enhanced the CS-US association in 60-s trace fear conditioning (Extended Data Fig. 6a & 6b). In contrast, chemogenetic activation of DRN-BA serotonergic neurons had no such effect (Extended Data Fig. 6c).

### Serotonin regulates vCA1 pyramidal neurons via 5-HT2C receptor during trace fear conditioning

To assess how serotonin modulates the vCA1 during trace fear conditioning, we used a recently developed multiplex fluorescence *in situ* hybridization (FISH) technique, PRISM^36^, to identify the cell type and 5-HTR expression patterns of neurons activated during training (Fig. 4a & 4b). Following 10-s trace fear conditioning, we sacrificed the mice and processed brain slices containing the vCA1 for RNA probing. Activated neurons were identified based on their expression of the immediate early gene *Fos*^37^. Based on previously reported RNA sequencing data in hippocampal neurons^38,39^, we probed the following five marker genes to classify the neuronal subtypes in the vCA1: *Slc17a7* (encoding vesicular glutamate transporter 1), *Gad1* (encoding glutamate decarboxylase 1, the synthetic enzyme for γ-aminobutyric acid), *Sst*, *Pvalb*, and *Vip* (encoding somatostatin, parvalbumin, and vasoactive intestinal peptide, respectively). In addition, we probed the genes encoding 5-HTRs previously implicated in fear conditioning, including *Htr1a*, *Htr1b*, *Htr2a*, *Htr2b*, *Htr2c*, and *Htr3a* (Fig. 4c & Extended Data Fig. 7a). Gene expression analysis revealed that the majority (∼80%) of *Fos*^+^ cells in the vCA1 were excitatory, with 30% of these cells expressing *Htr2c* (encoding Gq-coupled 5-HT2C receptor) and ∼18% expressing *Htr1a* (encoding Gi- coupled 5-HT1A receptor); only small percentages of *Fos*^+^/*Slc17a7*^+^ neurons expressed other 5-HTR subtypes (Fig. 4d & 4e). Consistent with these results, *ex vivo* imaging experiments showed that applying serotonin to acute brain slices induced calcium signal in ∼30% of vCA1 pyramidal neurons, which could be blocked by the selective 5-HT2C receptor antagonist, SB242084^(ref.40)^ (Extended Data Fig. 8 a-g).

**Fig. 4.**
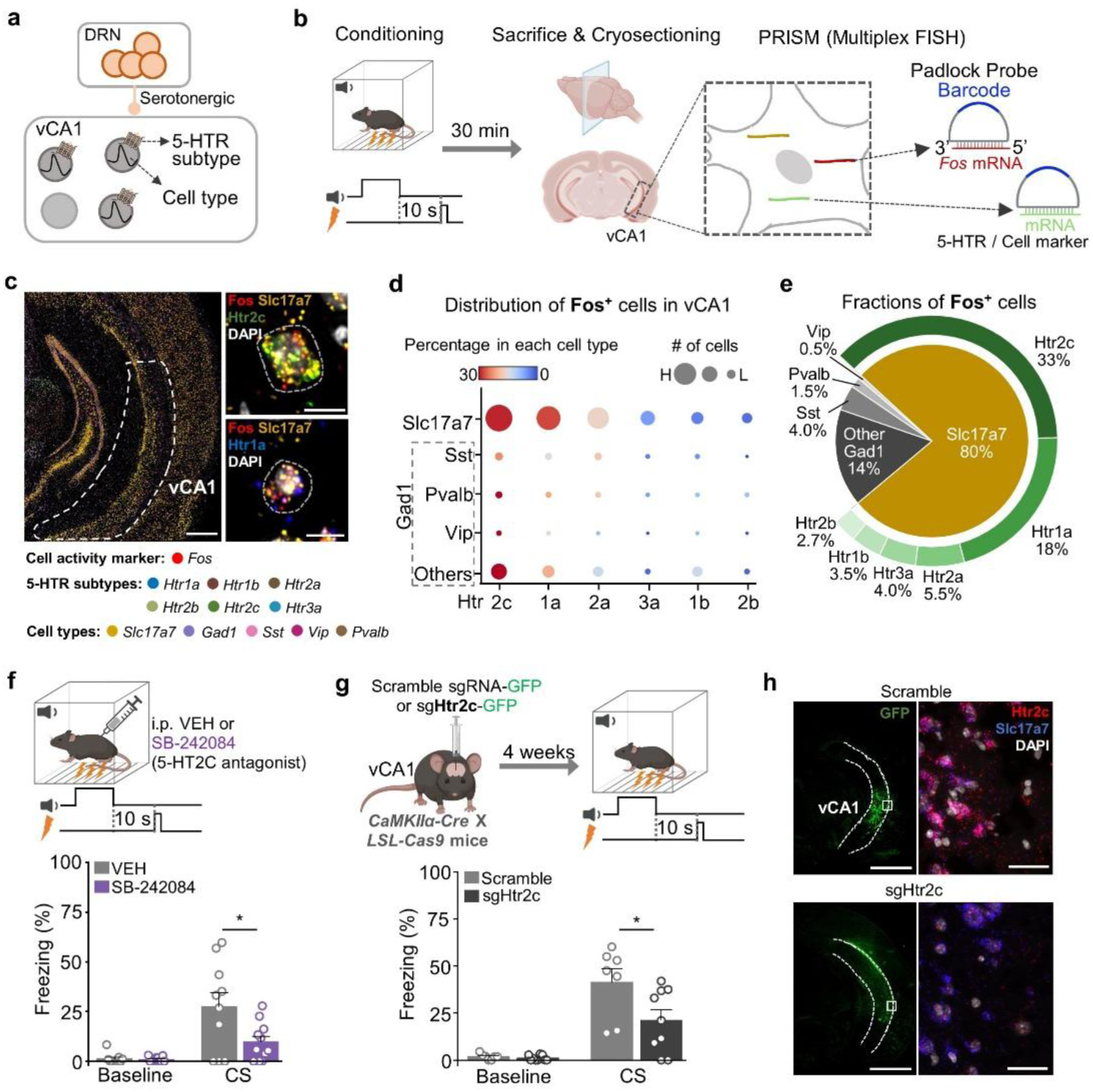
Serotonin regulates trace fear conditioning via 5-HT2C receptors in vCA1 pyramidal neurons. a &. **b.** Schematic of PRISM (multiplex FISH) on vCA1- containing brain slices after 10-s interval conditioning. Padlock probes targeted *fos*, 5- HTR genes, and cell type markers. **c.** PRISM showing spatial expression of *fos*, six 5- HTR genes, and five marker genes in the vCA1. High magnification insets (right) of selected genes are shown. Scale bars, 1 mm (left) and 10 μm (right). **d.** Dot plot of Fos^+^ cells distribution in the vCA1. **e.** Pie chart summarizing the fractions of Fos^+^ cells in the vCA1. **f.** Top: schematic of 10-s interval conditioning in mice treated with the 5- HT2C receptor antagonist SB-242084 (1 mg/kg) or vehicle (VEH). Bottom: summary of freezing time measured in the VEH and SB-242084 groups. * *p*=0.0281; unpaired *t* test; n=10-11 mice/group. **g.** Top: schematic of 10-s interval conditioning in *CaMKIIα- Cas9* mice injected with sgHtr2C (to knockdown 5-HT2C) or a scramble sgRNA. Bottom: summary of freezing time measured in indicated groups. * *p*=0.0387; unpaired *t* test; n=7-9 mice/group. **h.** Representative images of sgRNA expression in the vCA1 (left) and PRISM with *Htr2c* and *Slc17a7* probes to confirm *Htr2c* knockdown efficiency (right). Scale bars, 1 mm (left) and 100 μm (right). Data shown as means ± SEM.

To determine whether 5-HT2C receptors are critical for trace fear conditioning, we initially used pharmacological approaches. We found that an intraperitoneal (i.p.) injection of either SB242084 or the 5-HT2 type receptor antagonist Metergoline^41^ before 10-s trace fear conditioning significantly reduced freezing behavior compared to vehicle-treated control mice (Fig. 4f & Extended Data Fig. 7b). Given that ∼18% of trace fear conditioning-active neurons express the Gi-coupled 5-HT1A receptor, we also tested this receptor’s role and found that mice treated with 5-HT1A antagonist WAY- 100635^(ref.42)^ showed no change in freezing after 60-s trace fear conditioning compared to control mice (Extended Data Fig. 7c). Since the vCA1 neurons activated during training appear to constitute a molecularly defined cell type, we explored the effect of knocking out (KO) 5-HT2C receptors selectively in vCA1 pyramidal neurons using CRISPR-Cas9 editing^43^. We therefore designed an AAV construct containing five *Htr2c* sgRNA and a control AAV construct carrying a non-targeting scramble sgRNA (Extended Data Fig. 7d). We injected these AAVs into the vCA1 of *CaMKIIα/Cas9* mice, which express Cas9 specifically in excitatory neurons. We found that mice with conditional KO of *Htr2c* had significantly reduced freezing behavior compared to mice expressing the control AAV (Fig. 4g), confirming the role of 5-HT2C receptors in trace fear conditioning. The KO efficiency of *Htr2c* was confirmed by performing PRISM with specific probes targeting *Slc17a7* and *Htr2c* (Fig. 4h).

### Serotonin regulates vCA1 pyramidal neuron ensembles during trace fear conditioning through 5-HT2C receptor

We then examined how US-evoked serotonin release regulates vCA1 pyramidal cells in promoting the CS-US association. We initially recorded calcium activity in vCA1 pyramidal neurons during both the conditioning and the testing. We expressed the calcium sensor GCaMP6f^44^ under the *CaMKIIα* promoter in the vCA1 to image bulk calcium signals from pyramidal neurons using fiber photometry (Fig. 5a). In this experiment, we only applied pharmacological treatments before the conditioning. During 10-s trace conditioning, we observed a US-induced calcium signal, which was significantly reduced (in terms of both the peak response and the AUC) in mice pre-treated with SB242084 to block 5-HT2C receptors (Fig. 5b). These results highlight the critical role played by serotonin and the 5-HT2C receptor in regulating vCA1 neuronal activities during trace fear conditioning. After training, the CS-induced calcium signals increased, and were highly correlated with freezing behavior measured during the test. In contrast, SB242084-treated mice showed no memory retrieval, as evidenced by a lack of both calcium signals and freezing behavior in response to the CS (Fig. 5c-e). In a separate cohort, we examined the effect of fluoxetine on neuronal ensemble activity during 60-s trace conditioning and found that the US-induced calcium signal was transient, similar to serotonin release pattern. However, fluoxetine treatment significantly enhanced this calcium signal compared to vehicle treatment, indicating that increased serotonin levels promote the group activity of vCA1 pyramidal neurons (Fig. 5f). Moreover, consistent with fluoxetine-increased freezing behavior, we observed larger CS-induced calcium signals in the vCA1 of fluoxetine-treated mice compared to vehicle-treated ones (Fig. 5g-i).

**Fig. 5.**
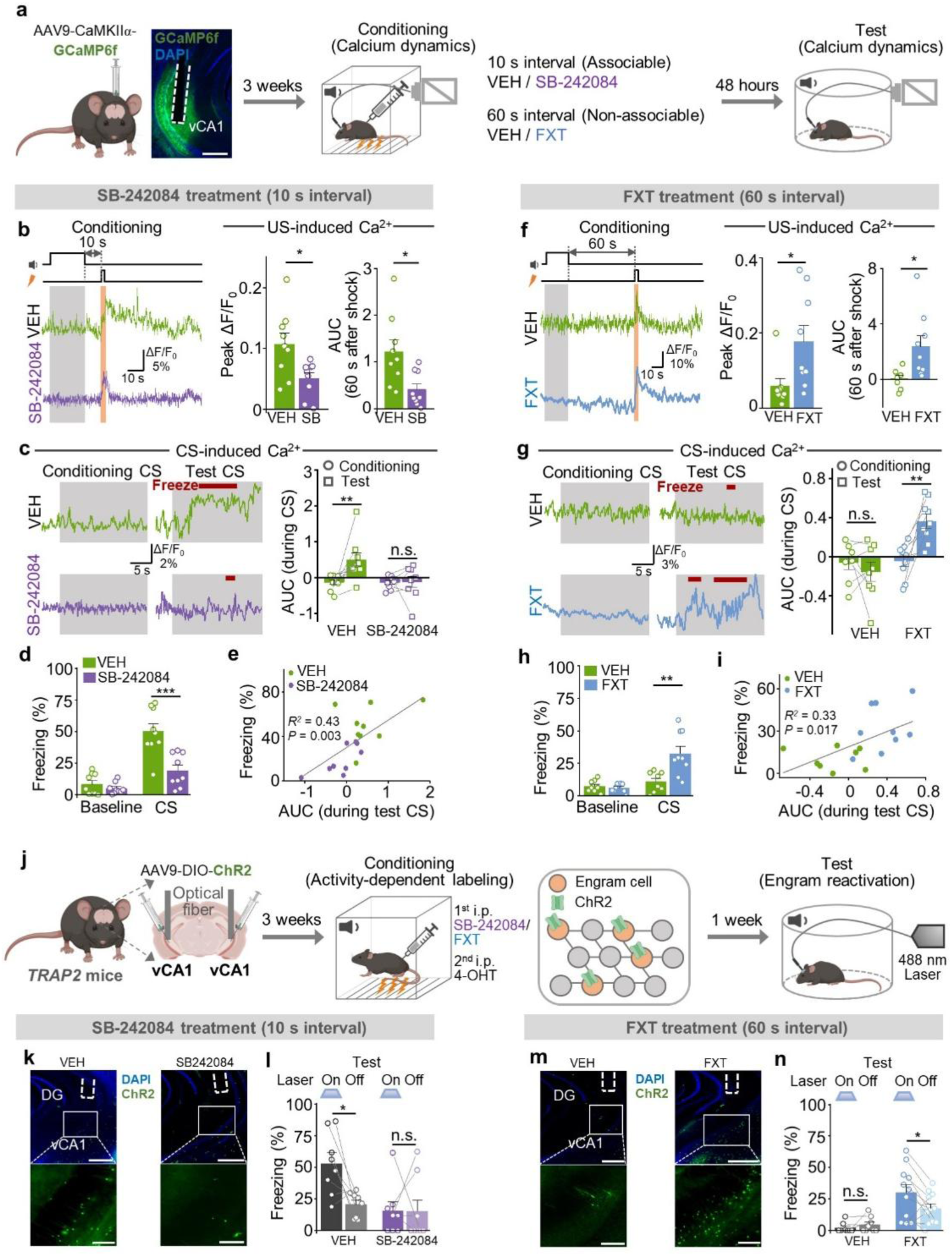
Serotonin regulates vCA1 pyramidal neuron ensembles during associable trace fear conditioning. **a.** Fiber photometry recordings in WT mice expressing GCaMP6f under the *CaMKIIα* promoter (excitatory neurons). Scale bar, 500 μm. **b.** Left: example traces (ΔF/F0) of GCaMP6f-expressing neurons during 10-s interval conditioning in mice treated with VEH or SB-242084 (1 mg/kg). Right: summary of peak ΔF/F0 and AUC of US-induced GCaMP6f signals. * *p*=0.0159 (peak), * *p*=0.0110 (AUC); unpaired *t* test; n=9 mice/group. **c.** Left: example traces during CS presentation in the conditioning (left) and test (right) sessions. Right: summary of the GCaMP6f signals during conditioning CS and test CS in the VEH and SB-242084 groups. ** *p*=0.0093; paired *t* test. **d.** Summary of freezing time (%) in the VEH and SB-242084 groups. *** *p*=0.0008; unpaired *t* test. **e.** Correlation between freezing behavior and AUC of the GCaMP6f signals during the test CS. Pearson’s correlation coefficient is shown. **f.** Left: example traces (ΔF/F0) of GCaMP6f-expressing neurons during 60-s interval conditioning in mice treated with VEH or FXT. Right: summary of peak ΔF/F0 and AUC of US-induced GCaMP6f signals. * *p*=0.0274 (peak), * *p*=0.0172 (AUC); unpaired *t* test; n=8-9 mice/group. **g.** Left: example traces during CS presentation in the conditioning (left) and the test (right) sessions. Right: summary of the GCaMP6f signals during conditioning CS and test CS in the VEH and FXT groups. ** *p*=0.0037; paired *t* test. **h.** Summary of freezing time (%) in the VEH and FXT groups. ** *p*=0.0042; unpaired *t* test. **i.** Correlation between freezing behavior and AUC of the GCaMP6f signals during the test CS. Pearson’s correlation coefficient is shown. **j.** Activity-dependent labeling of vCA1 neurons with ChR2 in *TRAP2* mice. Mice received 4-OHT (50 mg/kg) before conditioning to enable labeling of the neurons activated during conditioning. SB-242084 (1 mg/kg) or FXT (10 mg/kg) was used to manipulate the serotonergic system during conditioning. Engram reactivation was tested using 488-nm light during memory retrieval. **k & m.** Example images of ChR2 expression in the vCA1 in the indicated groups. Scale bars, 500 μm (upper) and 200 μm (lower). **l & n.** Summary of freezing time during “laser on” vs. “laser off” sessions in 10-s interval conditioning (**l**) and 60-s interval conditioning (**n**) in the indicated groups. 10 s: * *p*=0.0226; paired *t* test; n=8 mice/group. 60 s: * *p*=0.0353; paired *t* test; n=8-11 mice/group. Data shown as means ± SEM.

It has been proposed that neurons activated during training are preferentially allocated to an engram^45^, which is then reactivated during the test to enable memory retrieval^46^. To test whether sustained serotonin release regulates neuronal ensembles via 5-HT2C receptors in the vCA1, we used immunohistochemistry to quantify engram reactivation (Extended Data Fig. 9a). During training, active neurons were tagged with tdTomato (TdT) using the TRAP2 system. We found that mice treated with the 5-HT2C receptor antagonist SB242084 had fewer tagged (i.e., TdT^+^) neurons compared to vehicle-treated mice (Extended Data Fig. 9b-e). To assess engram reactivation during the test, we used a c-Fos antibody to label cells activated during the test and quantified neurons that were double-positive for TdT and c-Fos (TdT^+^ / c-Fos^+^). We found that SB-242084 treatment significantly reduced engram ensemble reactivation, as indicated by a reduced proportion of TdT^+^ / c-Fos^+^ cells in the vCA1 compared to vehicle-treated mice (Extended Data Fig. 9b-e).

Finally, to directly assess the role of serotonin in engram formation, we tagged ensemble neurons with ChR2 using *TRAP2* mice. Application of 4-OHT during training induced ChR2 expression in ensemble neurons, allowing for their subsequent activation with blue light during memory retrieval^47^ (Fig. 5j). In the memory test, we found that activating ChR2^+^ neurons significantly increased freezing behavior under 10-s interval conditioning; however, SB-242084 treatment eliminated this blue light-induced change in freezing behavior (Fig. 5k & 5l). On the other hand, 60-s interval conditioning induced minimal freezing that was not affected by blue light activation; however, fluoxetine treatment significantly increased freezing behavior when laser was on compared to laser off (Fig. 5m & 5n).

## Discussion

Here, we identified a distinct serotonergic circuit that regulates associability between the CS and the US in trace fear conditioning. Sustained serotonin release in the vCA1 following US delivery activates ensembles of 5-HT2C receptor-expressing pyramidal neurons, enabling the association between the temporally separated CS and US. While when the trace interval exceeds the associable window, the transient serotonin release in the vCA1 fails to induce CS-US association, thereby preventing maladaptive learning (Fig. 6). These findings therefore reveal a unique role for serotonin in regulating the temporal precision of associative fear learning.

**Fig. 6.**
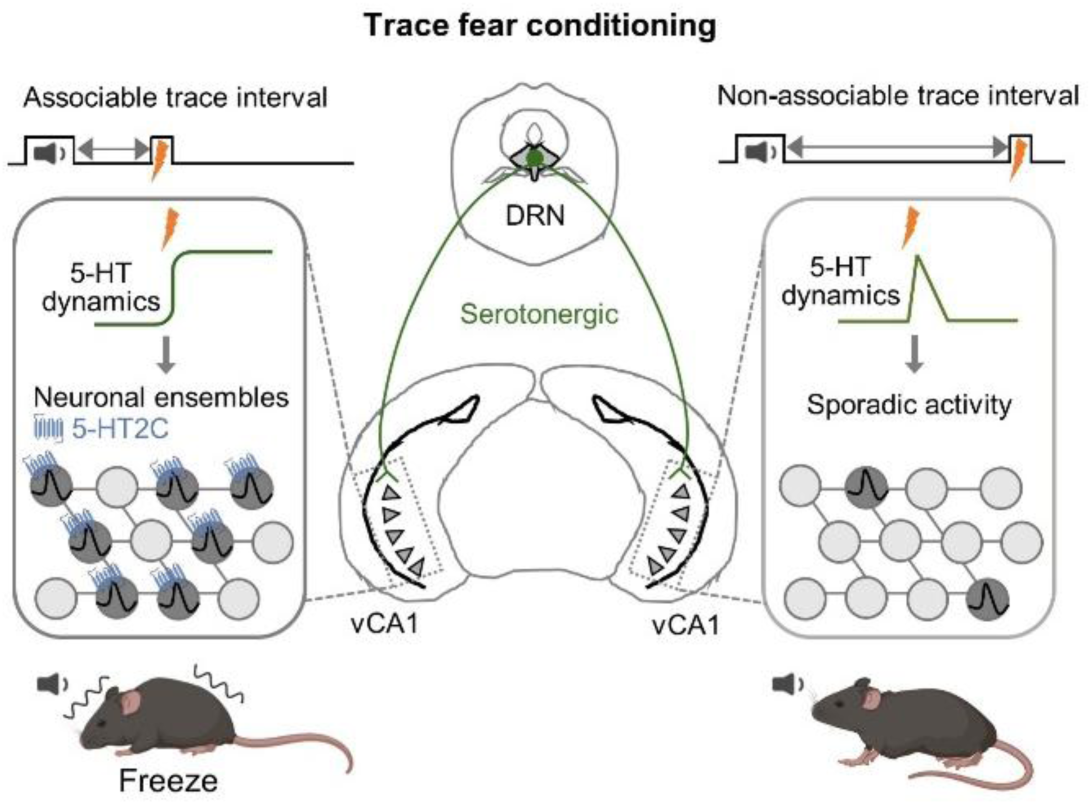
Working model. Schematic illustrating how the DRN-vCA1 serotonergic circuit regulates the associability of trace fear conditioning by actively promoting vCA1 neuronal ensembles.

The serotonergic system has been implicated in fear learning efficiency, particularly in delay fear conditioning^19,20^. Our work show that serotonin specifically modulates the associable window in trace fear conditioning via dynamic patterns of serotonin release in the vCA1. This serotonergic circuit, distinct from previously identified pathways, selectively regulates the association between discontiguous CS and US without affecting overall learning efficiency. We recently showed that serotonin regulates the CS-US association window in *Drosophila* aversive associative learning^48^, suggesting that this specific role of serotonin may be conserved across different species. Additionally, the 5-HT2C receptor has been implicated in fear conditioning. Pharmacological studies have shown that 5-HT2C receptor antagonists reduce the SSRI-induced increase in fear conditioning^18^, and knocking out the 5-HT2C receptor in mice attenuated fear conditioning^32,49^. Our results therefore provide a mechanistic link between serotonin signaling in the vCA1 and the temporal associability of fear conditioning, highlighting a novel role for serotonin and 5-HT2C receptor in regulating neuronal ensemble activity during trace fear conditioning.

Trace fear conditioning has been widely utilized to study the dynamic recruitment of brain regions involved in complex associative learning. Compared to delay fear conditioning, other brain regions in addition to the amygdala—including the hippocampus—have been implicated in CS-US association in trace fear conditioning. Lesion studies have shown that the dorsal hippocampus, particularly the dCA1, plays an essential role in trace fear conditioning^8^. However, in our study we observed minimal activity-dependent labeling and no detectable serotonin signal (data not shown) in the dCA1 during the training phase. Instead, we found that the vCA1 plays a critical role in trace fear conditioning. The hippocampus is functionally heterogenous along its dorsoventral axis, with the dorsal and ventral regions primarily involved in spatial memory and emotional memory, respectively^7^. This functional heterogeneity aligns with our observation that the vCA1 is required for trace fear conditioning, underscoring the role that the ventral hippocampus plays an important role in emotional memory. Recent studies have shown that vCA1 neuronal ensembles encode contextual fear memory^50,51^. Our results further suggest that a subset of vCA1 neuronal ensembles, regulated by serotonin, is essential for trace fear conditioning, providing new insights into the neural mechanisms underlying temporal associative learning.

Efficient and accurate fear learning is essential for survival, as this process ensures development of an adaptive responses while preventing maladaptive associations^52^. Thus, the regulation of temporal association is critical for fear learning. Studies regarding other forms of temporal associative learning have highlighted the hippocampus as a key brain structure that bridges temporally separated events^23,53^. Moreover, calcium imaging studies have shown that CS-induced neuronal activity in the hippocampus extends toward the US during trace fear conditioning^54^. However, recent evidence suggests that the hippocampus does not bridge the trace interval by generating persistent activity, but rather by altering its active neural population^55^. In our study, we shown that serotonin release in the vCA1 is precisely regulated to induce broad changes in pyramidal neuron activity, thus maintaining the associable time window. However, the brain regions responsible for regulating patterns of serotonin release under associable and non-associable trace intervals remain unclear. It is possible that the brain regions encoding the passage of time following the CS (auditory cue) regulate the DRN, thereby modulating serotonergic neuron activity based on the length of the interval between the CS and the US. Previous tracing studies have shown that serotonergic neurons in the DRN receive projections from the geniculate nucleus (GEN) of the thalamus^56^, a critical relay for all ascending auditory information to the cortex^57^ and an essential component of auditory fear conditioning neural circuit^58^. Given these anatomical connections, it is plausible that GEN inputs contribute to DRN serotonergic regulation. Future research is needed to elucidate the intrinsic mechanisms underlying this precise regulation during trace fear conditioning. Finally, our findings have important clinical implications, as altered regulation of serotonin signaling and maladaptive fear associations are hallmarks of anxiety-related disorders. Thus, targeting the DRN-vCA1 serotonergic pathway may provide new therapeutic strategies for these conditions.

## Methods

### Subjects

All experimental procedures involving animals were conducted in accordance with protocols approved by the Animal Care and Use Committee at Peking University. Male or female C57BL/6J mice (8-10 weeks old) were used for the experiments. Mice were pair-housed in a temperature-controlled (18–23 °C) and humidity-controlled (40–60%) room under a 12-h light /12-h dark cycle (light from 7 a.m. to 7 p.m.), with ad libitum access to food and water. All experiments were performed during the light cycle.

The following mouse lines were used for activity-dependent labeling, optogenetic, chemogenetic, and fiber photometry experiments: Sert-Cre mice (JAX strain 014554), TRAP2 mice (JAX strain 030323), Ai14 mice (JAX strain 007914), CaMKIIα-Cre mice (JAX strain 005359), Rosa 26-CAG-LSL-Cas9-tdTomato (generated by GemPharmatech). Male TRAP2xAi14 mice were bred in-house by crossing TRAP2 and Ai14 lines, and male CaMKIIα-Cas9 mice were bred in-house by crossing CaMKIIα-Cre and Rosa 26-CAG-LSL-Cas9-tdTomato lines. Only male mice were used for these experiments. The Sert-Cre mice were generously provided by M. Luo at Chinese Institute for Brain Research. The TRAP2 and the Ai14 mice were generously provided by C. Miao at Peking University.

### Viral vectors

For measuring serotonin dynamics, we used AAV9-hSyn-g5-HT3.0 (7.35x10^13^ vg/ml, Vigene Biosciences). For optogenetic experiments, we used AAV2/9-EF1α-DIO- hChR2(H134R)-mCherry (5.36x10^12^ vg/ml, BrainVTA), AAV2/9-EF1α-DIO- eNpHR3.0-mCherry (5.24x10^12^ vg/ml, BrainVTA), and AAV2/9-EF1α-DIO-mCherry (2.91x10^12^ vg/ml, BrainVTA). For chemogenetic experiments, we used AAV2/Retro- EF1α-DIO-FLP (5.93x10^12^ vg/ml, BrainVTA), AAV2/9-hSyn-fDIO-hM3D(Gq)-mCherry (2.73x10^12^ vg/ml, BrainVTA), and AAV2/9-hSyn-fDIO-mCherry (2.46x10^12^ vg/ml, BrainVTA). For conditional knockdown of 5-HT2C, we used AAV9-5-HT2C sgRNA (9.02x10^13^ vg/ml, Vigene Biosciences) and AAV9-scramble sgRNA (7.92x10^13^ vg/ml, Vigene Biosciences), that were driven by U6 promoter. For *in vivo* calcium recording, we used AAV2/9-CaMKIIα-GCaMP6f (2.08x10^12^ vg/ml, BrainVTA). For *ex vivo* calcium imaging, we used AAV2/9-CaMKIIα-GCaMP6s (2.35x10^12^ vg/ml, BrainVTA).

### Behavioral assays

All mice were handled daily for at least 3 days prior to behavioral experiment. On the day of testing, the mice were acclimated to the test room for a minimum of 1 h before experimentation. All behavioral procedures were performed during the light cycle and scored by the experimenters blinded to the treatment conditions.

#### Trace fear conditioning paradigm

Prior to all experiments, mice were handled for 1-min daily in the vivarium for 5 days. A four-day protocol was used for cued trace fear conditioning. On day 1 (habituation), mice were placed into a fear conditioning chamber (Med Associates) with a grid floor for 20-min to acclimate to the context. On day 2 (conditioning), after a 2-min baseline period, mice were exposed to a 20-s tone (5 kHz, 80 dB) followed by a trace interval and a 2-s scrambled foot shock (0.5 mA). A single conditioning trial was used to minimize the impact of repeated training on freezing behavior. Mice remained in the chamber for a 1-min consolidation period post-shock. For pharmacological experiments, drugs were administered via intraperitoneally (i.p.) injection 45 – 60 min before the conditioning session. To assess the effect of fluoxetine on memory consolidation, mice received an i.p. injection of fluoxetine within 1 h after conditioning. For delay conditioning experiments, mice were exposed to a 20-s tone (5 kHz, 80 dB) that co-terminated with a 2-s scrambled foot shock (0.5 mA). On day 3 (context extinction), mice were placed into the conditioning chamber for a 10 min context extinction session to eliminate context-dependent fear learning. On day 4 (test), mice were placed in a distinct conditioning chamber (Med Associates) featuring a white Plexiglas floor, dim lighting, a curved wall or A-frame insert, and was cleaned and scented with 1% acetic acid. After a 2-min baseline period, mice were presented with the a 20-s tone (5 kHz, 80 dB). Freezing behavior, defined as the cessation of all movement except for respiration, was quantified using an automated scoring system (Med Associates) with a sampling rate of 30 frames per second. In experiments involving tethered fibers, freezing was analyzed using a previously published open-source behavioral tracking pipeline, ezTrack^59^. Freezing was only counted if the mice remained immobile for at least 1 s.

#### Open-field test

The open-field test was conducted using ENV-510 test chambers (27.3 cm × 27.3 cm × 20.3 cm, Med Associates) equipped with infrared photo beams to assess spontaneous locomotor activity. At the start of the test, the mouse was placed in the center of the arena and allowed to freely explore the environment for 10-min. The mouse’s movement was tracked and analyzed using Activity Monitor 7 software (Med Associates) to measure the total travel distance and the time spent in the central zone (14.29 cm × 14.29 cm).

#### Elevated plus maze

The elevated plus maze consisted of two open arms (30 cm × 5 cm) without walls and two closed arms (30 cm × 5 cm × 15 cm) with opaque walls, connected by a central platform (5 cm × 5 cm), elevated 50 cm above the floor. At the beginning of each session, the mouse was placed in the center zone facing an open arm and allowed to explore the maze for 5 min. Locomotion trajectories were recorded using a video camera. The arena was thoroughly cleaned with 75% ethanol between trials. The total travel distance and the time spent in the open arms were quantified and analyzed using ezTrack.

### Drug Preparation

4-hydroxytamoxifen (4-OHT; Sigma, H6278) was prepared as previously described^27^. Briefly, 4-OHT was dissolved at 20 mg/ml in ethanol by shaking at 37°C for 15 min, followed by the addition of a 1:4 mixture of castor oil (Bide Pharmatch, BD01411804) and sunflower seed oil (MREDA, M109741) to achieve a final concentration of 10 mg/ml 4-OHT. Ethanol was evaporated under vacuum centrifugation, resulting a final working solution of 20 mg/ml. The working solution was prepared fresh on the day of use and administered i.p. at a dose of 50 mg/kg.

Fluoxetine hydrochloride (FXT; Sigma, F132) was dissolved at 1 mg/ml in saline and delivered i.p. at a dose of 10 mg/kg. SB-242084 (Abmole, M3772) was dissolved at 100 μg/ml in 1% DMSO and delivered i.p. at a dose of 1 mg/kg. WAY-100635 maleate salt (Sigma, W108) was dissolved at 100 μg/ml in 1% DMSO and delivered i.p. at a dose of 1 mg/kg. Metergoline (Sigma, M3668) was dissolved at 200 μg/ml in 1% DMSO and delivered i.p. at a dose of 2 mg/kg. Deschloroclozapine (DCZ; MCE, HY-42110) was dissolved at 10 μg/ml in 1% DMSO and delivered i.p. at a dose of 100 μg/kg.

### Activity-dependent labeling and optogenetic manipulation

We used *TRAP2 x Ai14* mice^27^ to achieve activity-dependent labeling of vCA1 neurons in response to trace fear conditioning. Neuronal activation induces the expression of CreERT2 recombinase, which translocates to the nucleus upon 4-hydroxytamoxifen (4- OHT) injection and catalyzes recombination, leading to permanent tdTomato expression. The mice were habituated to the conditioning chamber for 20 min daily and received a mock injection before each habituation session for 1 week to minimize non-specific c-Fos induction. After the animals were fully habituated to the test room, they received an i.p. injection of 4-OHT (50 mg/kg), followed 30 min later by either 10-s or 60-s interval trace fear conditioning, or exposure to the conditioning chamber alone. Mice were then returned to their home cages and were sacrificed 1 week after TRAPing.

The activity-dependent labeling of vCA1 cells was validated by assessing the overlap between trace conditioning-TRAPed cells and cells expressing c-Fos. In brief, 90 min after the test session the mice were anesthetized and transcardially perfused with 4% paraformaldehyde (PFA).

For optogenetic manipulation of TRAPed cells, we first injected AAV-DIO-ChR2 into the bilateral vCA1 of TRAP2 mice. Three weeks after AAV injection, the mice underwent TRAP labeling as described above; One week later, each animal was subjected to a test session. During the test, the CS was delivered twice (20 s each), once in the presence and once in the absence of blue light in a counterbalanced order.

### Stereotaxic surgery and targeting coordinates

Anesthesia was induced at 3% isoflurane in oxygen (2 L/min) and maintained at 1–2%. For virus injection and fiber implantation, the mouse was secured in a stereotactic frame (RWD Instruments). The following stereotactic coordinates were used for the indicated brain regions: vCA1 (AP: -3.2 mm relative to bregma; ML: ±3.3 mm relative to bregma; DV: −3.4 mm from the dura), BA (AP: -1.35 mm relative to bregma; ML: ±3.0 mm relative to bregma; DV: -4.5 from the dura), PFC (AP: +1.7 mm relative to bregma; ML: ±0.3 mm relative to bregma; DV: −1.85 mm from the dura), DRN (AP: -4.3 mm relative to bregma; ML: ±1.1 mm relative to bregma (with a 20° angle); DV: −2.85 mm from the dura), and MRN (AP: -4.3 mm relative to bregma; ML: ±0.2 mm relative to bregma; DV: −4.15 mm from the dura). Viral solutions (300 nL) was injected either unilaterally or bilaterally into the target brain regions using a pulled glass capillary attached to a pressure microinjector (World Precision Instruments). The injection needle was kept in place for 5 min post-injection to minimize backflow. Optical fibers (200-μm diameter, 0.37 NA; Inper Ltd.) were implanted at the AAV injection site and secured with dental cement (3M). The optical fibers were placed either 0.15 mm or 0.4 mm above the virus injection sites for fiber photometry recording experiments and optogenetic experiments, respectively. After surgery, mice were allowed to recover from anaesthesia on a heating pad.

### Fiber photometry recording

Three weeks after virus injection, fiber photometry recordings were performed using an FPS-410/470 photometry system (Inper Ltd.) and signal v.2.0.0 software (Inper Ltd.). Briefly, a 470/5-nm filtered light-emitting diode (LED) at 20-30 μW (10 Hz, 20 ms pulse duration) and a 410/5-nm filtered LED at 20-30 μW (10 Hz) were used to excite 5-HT3.0 or GCaMP6f. Alternating excitation wavelengths were delivered, and the fluorescence signals were collected using a sCMOS camera during dual-color imaging. To minimize autofluorescence of the optical fiber, the recording fiber was photobleached using a high-power LED prior to recording. Photometry data were analyzed using a custom MATLAB script (MATLAB R2022a, MathWorks). Specifically, the 410-nm channel signal was aligned to the 470-nm channel signal by linear fitting. The 470-nm channel signal was then normalized to the fitted control by calculating the change in fluorescence [F = (470-nm channel signal) / (fitted 410-nm channel signal)]. To calculate ΔF/F0 during the conditioning session, baseline fluorescence (F0) was measured 5 s before the indicated stimuli (either CS or US delivery). For comparison of signal between animals, ΔF/F0 was further normalized, and the area under the curve (AUC) was calculated. To analyze the 5-HT3.0 signal during the test session, z score‒transformed signals were calculated using the standard deviation of the baseline signals. The code for fiber photometry recording analysis has been deposited on GitHub (https://github.com/yulonglilab/FiberPhotometry).

### Optogenetics

To activate the synaptic terminals, blue (470 nm) laser light was bilaterally delivered onto ChR2- expressing DRN serotonergic axon terminals in the vCA1. The laser power was adjusted to ∼10 mW at the tip of the optical fiber. Before conditioning each mouse, the laser power was calibrated by measuring the output at the tip of the optical fiber patch cord (Inper Ltd.). Optogenetic activation was delivered at 20 Hz (5 ms pulses) to mimic the previously reported firing rate of serotonergic neurons during exposure to aversive stimuli^60^. To inhibit activity at the synaptic terminals, yellow (556-nm) laser light was bilaterally delivered continuously (∼10 mW from the optic fiber tip) to eNpHR3.0- expressing DRN or MRN serotonergic axon terminals in the vCA1. Light delivery was terminated using a 10-second ramping down protocol to attenuate the rebound effect of eNpHR3.0^61^.

### Chemogenetics

For chemogenetic activation of the DRN-vCA1 or DRN-BA serotonergic pathway, AAVretro-DIO-FLP was injected bilateral into the vCA1 or BA, and AAV-fDIO-hM3Dq was injected into the DRN of *Sert-Cre* mice. The trace fear conditioning paradigm was conducted 3 weeks after virus injection. Thirty minutes before the conditioning session, the mice received an i.p. injection of deschloroclozapine (DCZ, 100 μg/kg) to activate hM3Dq.

### Immunohistochemistry

Mice were anaesthetized using Avertin and intracardially perfused with phosphate-buffered saline (PBS) followed by 4% PFA in PBS. The brains were dissected and post-fixed in 4% PFA at 4°C overnight. Coronal sections (40-μm thick) were obtained using a VT1200 vibratome (Leica). Brain slices were incubated in blocking solution (5% normal goat serum, 0.1% Triton X-100 and 2 mM MgCl^2^ in PBS) for 1 h at room temperature. The slices were then incubated overnight at 4 °C in AGT solution (0.5% normal goat serum, 0.1% Triton X-100 and 2 mM MgCl2 in PBS) containing the following primary antibodies: anti-5-HT antibody (ImmunoStar, 20080, rabbit, 1:1,000 dilution) or anti-c-Fos antibody (Abcam, AB222699, rabbit, 1:1,000 dilution). The following day, the slices were rinsed three times in AGT solution and incubated for 1 h at room temperature with the following secondary antibodies: Goat anti-rabbit Alexa Fluor 488 (Abcam, ab150169, 1:500 dilution) or Goat anti-Rabbit iFluor 555 (AAT Bioquest, 16690, 1:500 dilution). After three additional washes in AGT, the slices were mounted on slides using a mounting medium containing DAPI (Abmole BioScience, 0100-20) and sealed with coverslips. The slides were then imaged using a VS120-S6 slide scanner (Olympus) equipped with a ×10 objective.

For TRAPed cell counting, the brains were sectioned at 40 μm thickness, and every third slice was collected for analysis. Anatomical landmarks based on the Allen Brain Atlas were used to identify specific brain regions. Cells were counted and analyzed offline in a blinded manner using ImageJ and Qupath software.

### PRISM (Super-multiplex FISH)

PRISM was conducted as previously described^36^. We designed multiple padlock probes for each detected gene in order to improve detection sensitivity (8 probes per gene for 5-HT2C knockdown validation and 3 probes per gene for other genes). The fresh-frozen mouse brains were sectioned into 10-μm-thick slices using a CM1950 Cryostat (Leica) and mounted on Superfrost Plus glass slides (Epredia). Tissue sections were fixed in 4%

PFA at room temperature for 15-min, followed by permeabilization with 0.01% pepsin in 0.1 M HCl at 37 °C for 2-min. Samples were then dehydrated through a series of ethanol washes (10-min in 80% ethanol, followed by 2-min in 100% ethanol) and rehydrated with three-time PBST (0.05% Tween 20 in 1xPBS) washes. Samples were blocked with oligo-dT (100 nM oligo-dT, 50 mM KCl, 20% formamide, 20 μg/mL BSA, 20 μg/mL Yeast tRNA (Invitrogen, AM7119), and 1U/μL Ribolock RNase inhibitor (Thermo Scientific) in 1xAmpligase buffer (Lucigen)) for 10 min at room temperature. Gene-specific padlock probes were hybridized to the samples by incubating them in a hybridization mix (200 nM padlock probe for each, 50 mM KCl, 20% formamide, 20 μg/mL BSA, 20 μg/mL Yeast tRNA, and 1U/μL Ribolock RNase inhibitor in 1xAmpligase buffer) at 55 °C for 10 min and then 45 °C for 1 h 50 min. Ligation was performed by incubating samples in a ligation mix (2.5 U/μL SplintR ligase (NEB), 20 μg/mL BSA, and 1 U/μL Ribolock RNase inhibitor in 1xSplintR buffer) at 37 °C for 2 h to seal nicks in the padlock probes. Rolling circle amplification (RCA) was performed using 0.25 U/µl 1xPhi29 polymerase in Phi29 polymerase buffer (Thermo Scientific) with 250 μM dNTP (Thermo Scientific), 50 μM aminoallyl-dUTP (Thermo Scientific), 10% glycerol, 20 μg/mL BSA, and 300 nM RCA primer at 30 °C overnight. After RCA, samples were fixed with 10 μg/μL BS(PEG)9 (Thermo Scientific) in PBST for 15 min at room temperature. For imaging probe staining, samples were incubated with a mixture of fluorescence-labeled probes (each paired with an unlabeled probe at a predetermined ratio, final concentration of 120 nM per pair, 20% formamide, 2X SSC buffer) at 50 °C for 9 min, followed by 37 °C for 21 min. Samples were then stained with DAPI (5 μg/μL, Beyotime) for nuclear visualization. To image 10-µm brain sections, we collected a z-stack of 9 planes at ∼1 µm intervals for each tile, imaging was performed with an epifluorescence microscope.

### PRISM image analysis

Imaging processing and gene decoding were performed as previously described^36^. Briefly, images from multiple z-planes were stacked using an all-in-focus algorithm, followed by shade correction with CIDRE and inter-channel image registration (Fast Fourier Transform and Maximum Cross-Correlation). Color aberration was corrected by resizing images from different channels. Tiles were stitched using the Microscopy Image Stitching Tool. Signal spots were extracted from all four channels by identifying local maxima on tophat-filtered images for thin tissue sections. Spot signal coordinates across all channels were combined to generate 4-dimensional intensity information (from Ch1 to Ch4). Intensity values were normalized using the mean values of their respective channels. Spots with low sum-intensity values were filtered out. After that, we plotted all spots in a “color space” (x = (Ch1-Ch2)/(Ch1+Ch2+Ch3), y = 2*Ch3/(Ch1+Ch2+Ch3)-1, and z = Ch4/(Ch1+Ch2+Ch3)) and performed gene calling according to spots’ position in color space. Cell nuclei were segmented using adaptive thresholding and watershed algorithm of DAPI images, after which we assigned each called transcript to nearest nucleus centroid to generate an expression matrix for the cells. Cells were briefly classified by thresholding corresponding marker gene expression. For detailed PRISM decoding workflow and following analysis, the code can be found in Github (https://github.com/huanglab111/PRISM).

### Two-photon imaging in acute mouse brain slices

Three weeks after CaMKIIα-GCaMP6s virus injection, mice were anesthetized with Avertin and intracardially perfused with ice-cold oxygenated slicing buffer containing (in mM): 110 choline chloride, 2.5 KCl, 1.25 NaH2PO4, 25 NaHCO3, 7 MgCl2, 25 glucose and 0.5 CaCl2. The brain was rapidly removed and placed in chilled, oxygenated slicing buffer. Coronal slices (300 μm thick) containing the vCA1 region were sectioned using a VT1200 vibratome (Leica) in cold slicing buffer, and then transferred to oxygenated artificial cerebrospinal fluid (aCSF) containing (in mM): 125 NaCl, 2.5 KCl, 1 NaH2PO4, 25 NaHCO3, 1.3 MgCl2, 25 glucose and 2 CaCl2. The slices were then incubated at 34 °C for recovery. Two-photon imaging was performed using an Ultima Investigator two-photon microscope (Bruker) equipped with a 20X/1.00-NA objective (Olympus), an InSight X3 tunable laser (Spectra-Physics), and Prairie View 5.5 software (Bruker). A 920-nm laser was used to excite the GCaMP6s sensor, and fluorescence was collected using a 495–540-nm filter. Where indicated, drugs were perfused at a flow rate of 4 mL/min.

### Statistical analysis

Mice were randomly assigned to the treatment groups, and all behavioral experiments were performed in a blinded manner. Statistical analyses were conducted using GraphPad Prism software (v9). Unless otherwise indicated, summary data are reported as the mean ± standard error of the mean (s.e.m.). Data were excluded from the analysis if the virus expression or fiber implantation sites were determined to be outside the target brain region. All datasets were assumed to be normally distributed, and equal variances were formally tested. Differences between groups were analyzed using the two-tailed Student’s paired or unpaired *t* tests, or a one-way ANOVA, as appropriate. Statistical significance was set at *p* < 0.05.

## Acknowledgements

We thank Dr. Yi Rao, Dr. Minmin Luo and Dr. Chenglin Miao for providing transgenic mouse lines. We are grateful to Dr. Zachary Pennington for his instructive suggestions on the trace fear conditioning paradigm design. We also thank National Center for Protein Sciences at Peking University in Beijing, China, for providing the behavioral setups. Some diagrams were created using BioRender.com. We thank Yueyue Yu, and Yuehan Chen for their diligent maintenance of the mouse colony. We thank the members of the Li lab for their valuable suggestions and comments on the manuscript.

## Author contributions

Y.L. and Y.Z. conceived and designed the project. Y.Z., X.L., T.C., S.X., S.L., J.C., and D.W. performed experiments. Y.Z., X.L., T.C., and Y.W. analyzed data. F.D. provided viral constructs. Y.Z. and Y.L. prepared figures and wrote the manuscript. The other authors made valuable comments on drafts of the manuscript and figures. Y.H. supervised multiplexed FISH experiments. Y.L. supervised the entire project.

