## Supplemental figures for "Serotonin shapes the temporal window for associative fear learning"

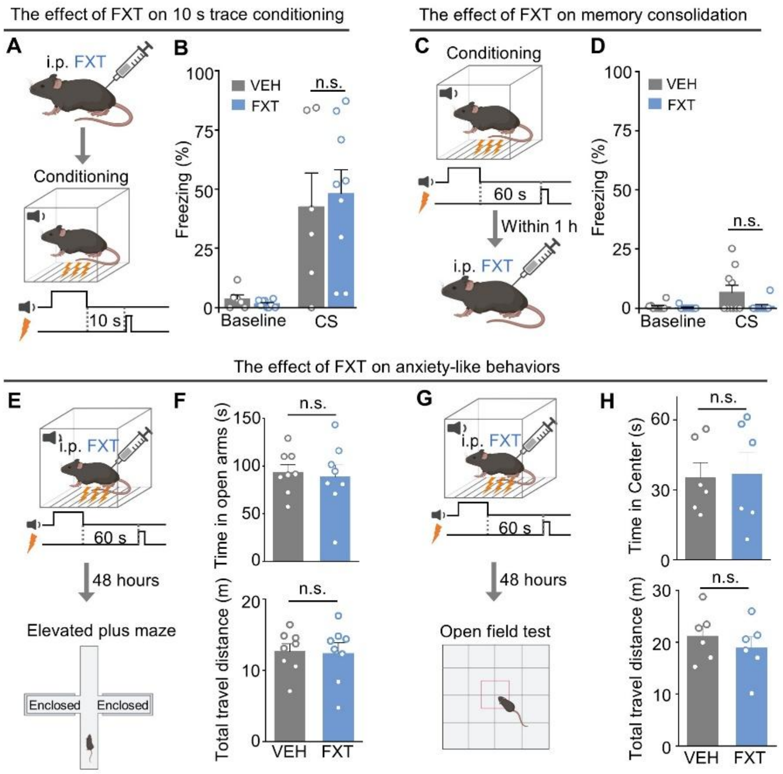


**Fig. S1. Effects of FXT injection on different behavioral tests.**

(**A** and **B**) Summary of freezing time measured in the VEH and FXT groups during 10-s interval conditioning. Unpaired *t* test; n=6-9 mice/group. (**C** and **D**) Summary of freezing time measured in the VEH and FXT groups when FXT was administrated post-conditioning. Unpaired *t* test; n=10 mice/group. (**E** to **H**) Anxiety-like behaviors in the FXT and VEH groups. Elevated plus maze test (E and F), unpaired *t* test; n=8 mice/group. Open field test (G and H), unpaired *t* test; n=6 mice/group. Data shown as means ± SEM.

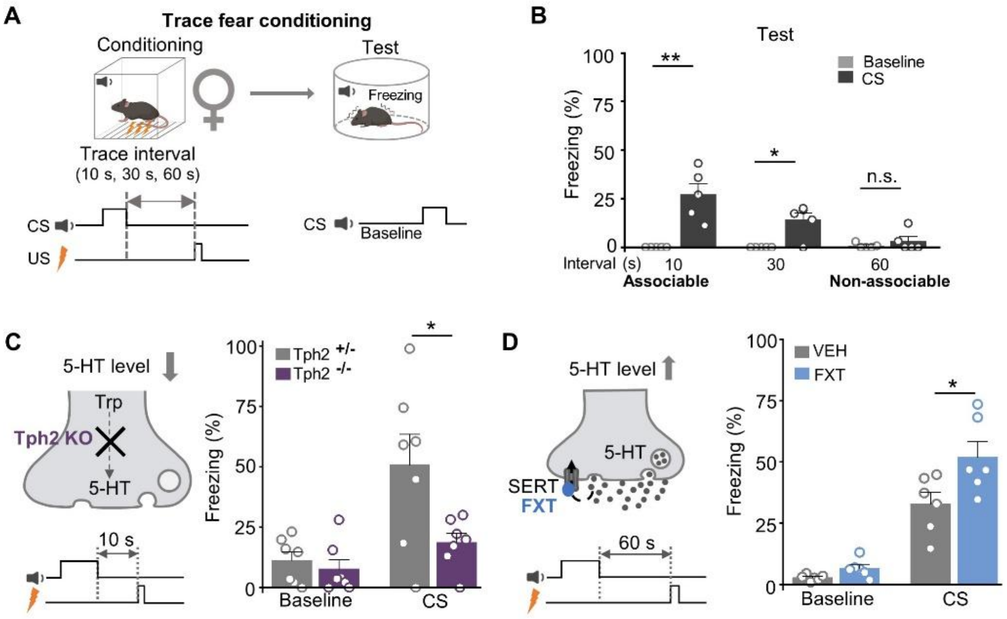


**Fig. S2. Behavioral changes of female mice upon serotonin system manipulations.** (**A** and **B**) Freezing time (%) during baseline and CS presentation in female mice. Baseline vs. CS, ** $p$=0.0098 (10 s), * $p$=0.0184 (30 s); paired $t$ test; n=5 mice/group. (**C**) Freezing time (%) of female *Tph2* KO mice during 10-s interval conditioning. * $p$=0.0310; unpaired $t$ test; n=7 mice/group. (**D**) Freezing time (%) of female mice treated with 10 mg/kg FXT during 60-s interval conditioning. * $p$=0.0356; unpaired $t$ test; n=6 mice/group.

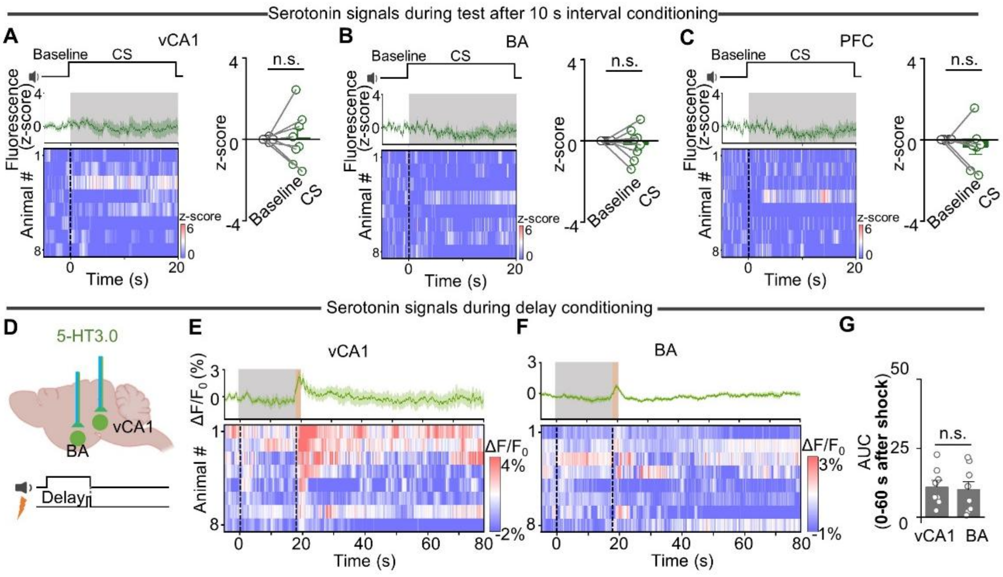


**Fig. S3. Fiber photometry recordings of serotonin dynamics.**

(**A** to **C**) Traces (left) and summary (right) of 5-HT3.0 signals during test sessions after 10-s interval conditioning in the vCA1 (A), BA (B), and PFC (C) (related to Fig. 2C). The grey shaded areas indicate the CS delivery. No apparent signals were measurable during CS presentation. Paired *t* test; n=8 mice/group. (**D** to **G**) 5-HT3.0 signals measured in the vCA1 and BA during delay fear conditioning. The grey and orange shaded areas indicate the CS and US delivery, respectively. Unpaired *t* test; n=8 mice/group. Data shown as means ± SEM.

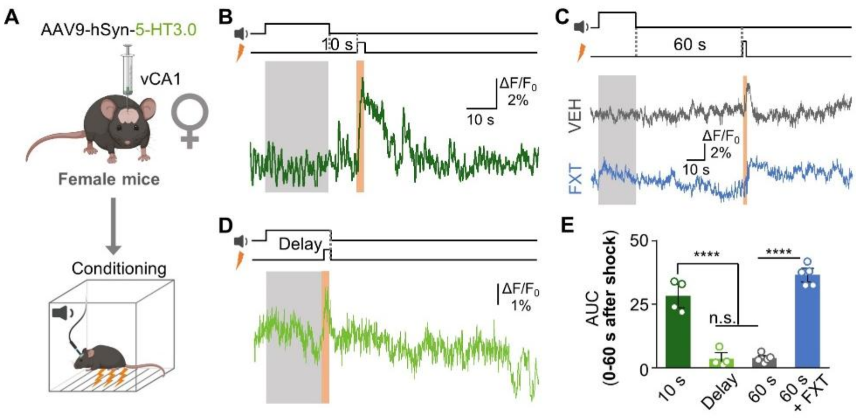


**Fig. S4. 5-HT3.0 signal patterns of female mice.**

(**A** to **D**) 5-HT3.0 signals measured in the vCA1 during 10-s (B), 60-s with or without FXT treatment (C) trace fear conditioning, and during delay fear conditioning (D). (**E**) Summary of AUC of the 5-HT3.0 signals. **** $p<0.0001$, One-way ANOVA with Tukey's post hoc test; n=4-5 mice/group. Data shown as means ± SEM.

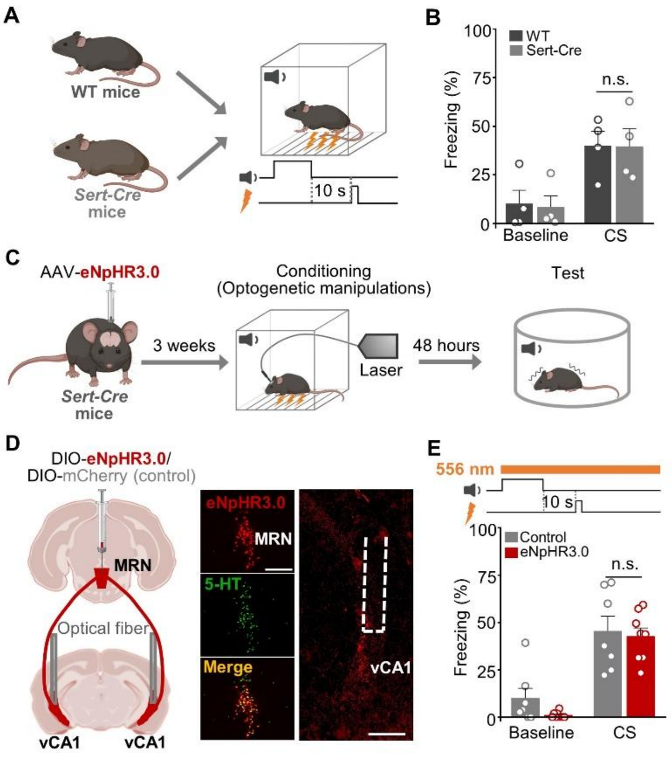


**Fig. S5. Optogenetic inhibition of MRN-vCA1 serotonergic projections has no effect on trace fear conditioning.**

(**A** and **B**) Summary of freezing time measured in WT and *Sert-Cre* mice. Unpaired *t* test; n=4 mice/group. (**C**) Timeline of the behavioral paradigm with optogenetic manipulation. (**D**) Left: graphical representation of AAV-DIO-eNpHR3.0 or AAV-DIO-mCherry injection into the DRN of *Sert-Cre* mice, with fiber implantation in the bilateral vCA1. Right: example images of eNpHR3.0 and 5-HT colocalization in the DRN, and eNpHR3.0 expression in the vCA1. Scale bar, 500 μm. (**E**) Summary of freezing time measured after optogenetic inhibition of the MRN-vCA1 serotonergic circuit during the entire conditioning session. Unpaired *t* test; n=7-8 mice/group. Data shown as means ± SEM.

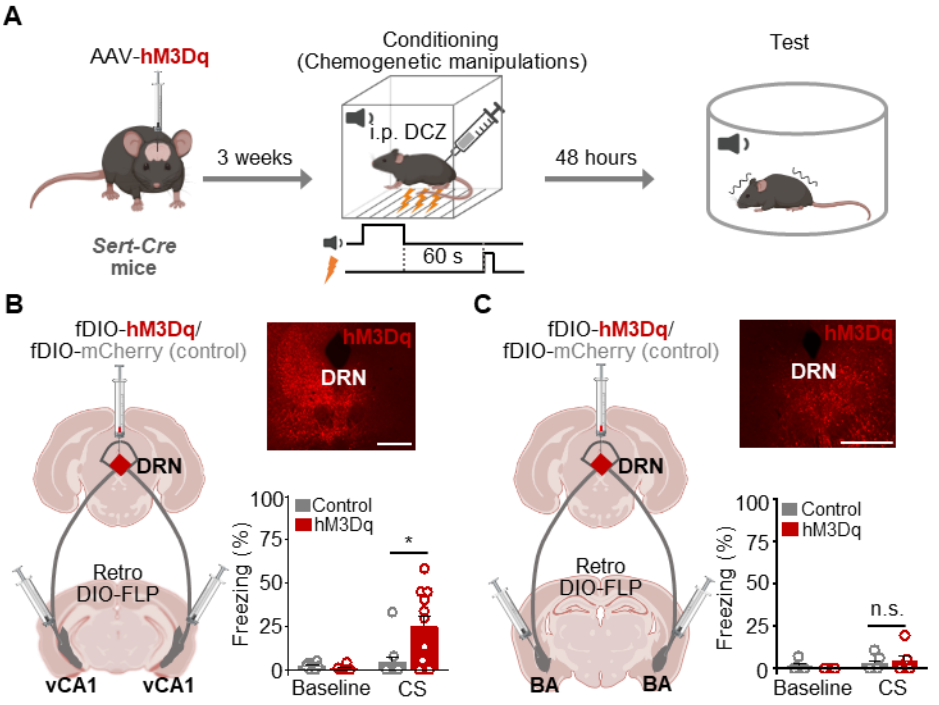


**Fig. S6. The effect of chemogenetic activation of different serotonergic circuits on non-associable trace fear conditioning.**

(**A**) Timeline of the behavioral paradigm with chemogenetic manipulation. The mice received 100 μg/kg Deschloroclozapine (DCZ) before conditioning to activate hM3Dq. (**B** and **C**) Left: graphical representation of AAV-fDIO-hM3Dq or AAV-fDIO-mCherry injection into the DRN, and AAV-DIO-FLP injection into the bilateral vCA1 (B) or BA (C) of Sert-Cre mice. Right: summary of freezing time measured in the control and hM3Dq groups following manipulation of the DRN-vCA1 serotonergic circuit (B) or the DRN-BA serotonergic circuit (C). * p=0.0135; unpaired t test; n=6-11 mice/group. Data shown as means ± SEM.

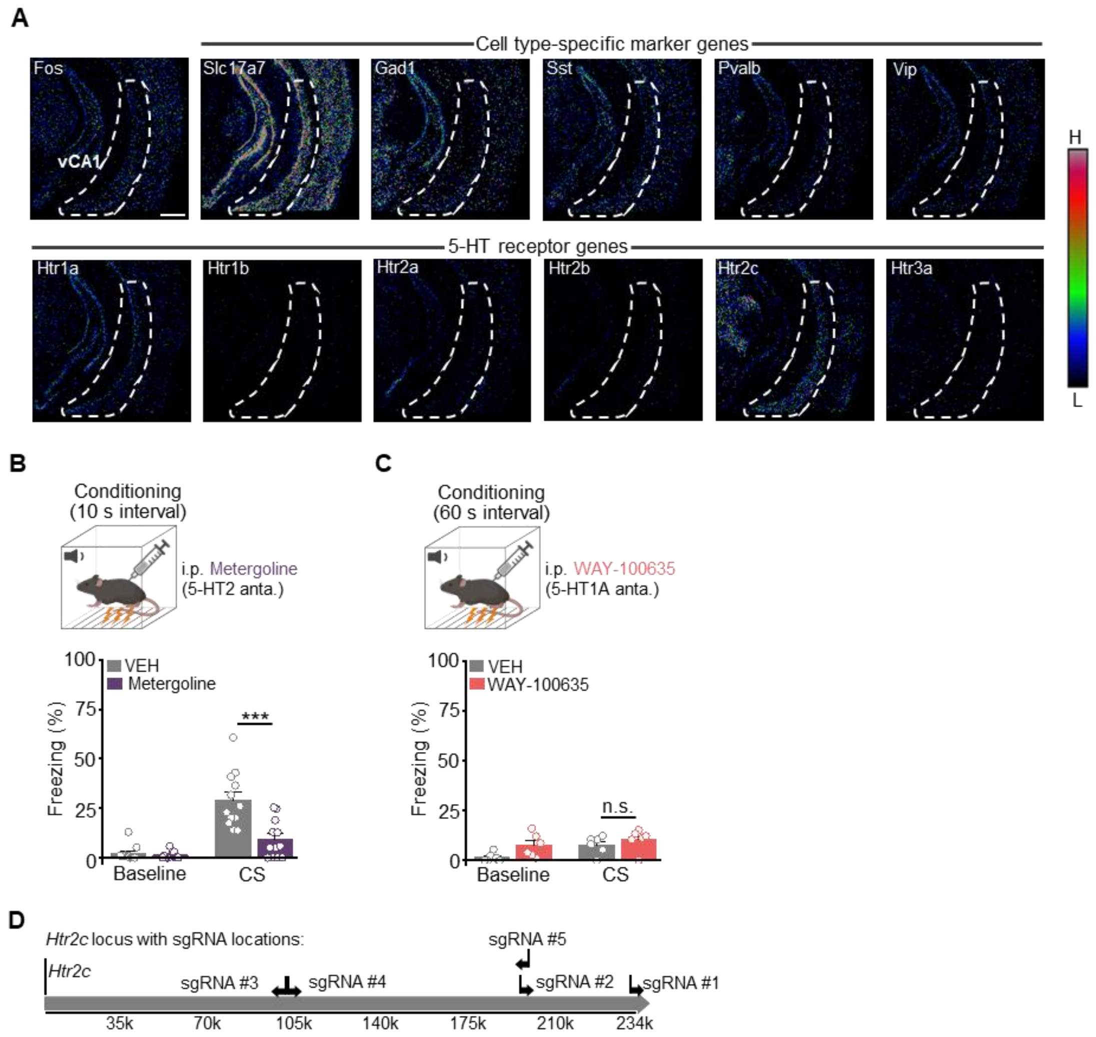


**Fig. S7. Pharmacological manipulation of 5-HT receptors.**

(**A**) FISH images showing *fos*, six 5-HTR genes, and five marker genes in vCA1 (related to Fig. 4B). (**B**) Freezing time measured in mice treated with 5-HT2 antagonist Metergoline (2 mg/kg) or VEH during 10-s interval conditioning. *** $p$=0.0006; unpaired $t$ test; n=12 mice/group. (**C**) Freezing time measured in mice treated with 5-HT1A antagonist WAY-100635 (1 mg/kg) or VEH during 60-s interval conditioning. Unpaired $t$ test; n=6 mice/group. (**D**) The *Htr2c* locus and constructs used for knocking down the 5-HT2C receptor. A mixture of 5 sgRNAs were used to make sgHtr2c and control construct expresses a non-targeting sgRNA. Data shown as means ± SEM.

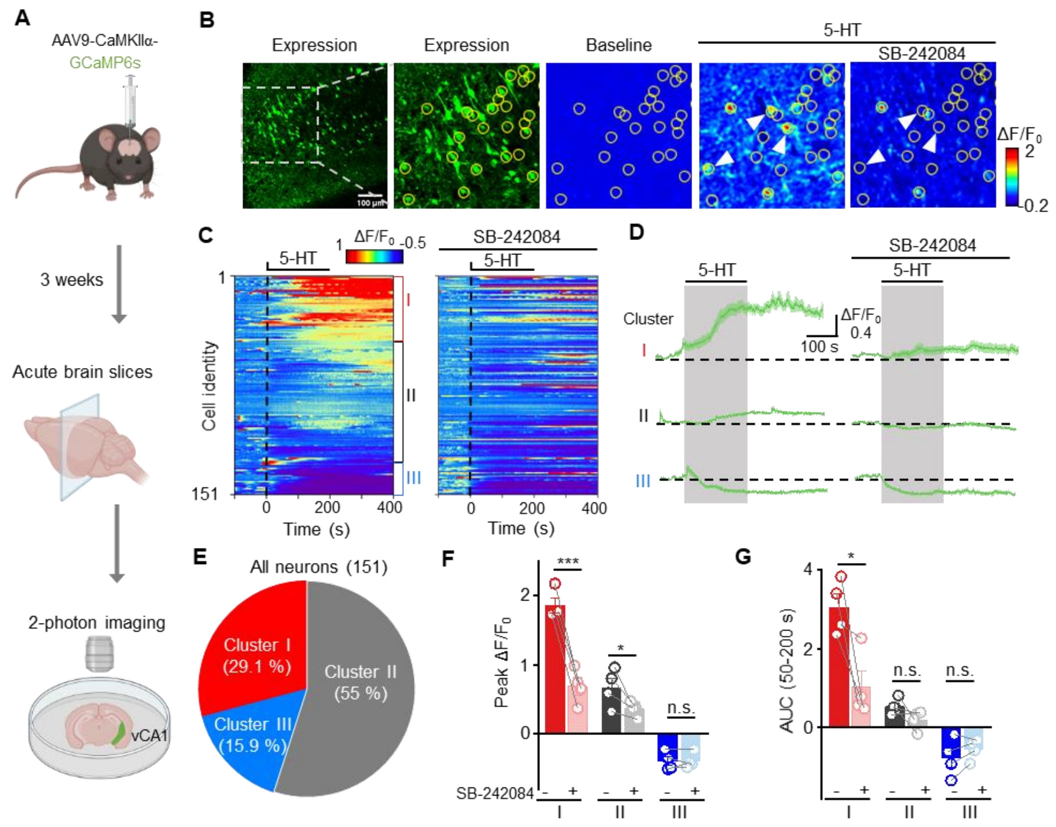


**Fig. S8. Serotonin induces calcium signals in vCA1 pyramidal neurons that can be blocked by 5-HT2C receptor antagonist.**

(**A**) Experimental design: AAV9-CaMKIIα-GCaMP6s was expressed in the vCA1 for 2-photon imaging of acute brain slices. (**B**) Left: two-photon images of GCaMP6s expression. Right: pseudocolor images at baseline and in response to 5-HT (10 μM) application with or without the 5-HT2C antagonist SB-242084 (10 μM). White arrows indicate cells with 5-HT-induced calcium signals that were blocked by SB-242084. (**C**) Example heatmap of calcium activity measured in vCA1 pyramidal cells, grouped by increased (Cluster I), unchanged (Cluster II), or decreased (Cluster III) activity in response to 5-HT. (**D**) Average calcium activity of the three cell clusters in the vCA1 of one brain slice. (**E**) Pie chart showing the distribution of cells in each cluster. (**F** and **G**) Summary of peak $\Delta F/F_0$ (F) and AUC (G) of the calcium signals in each cluster. For peak $\Delta F/F_0$, *** $p$=0.0005, * $p$=0.0389; For AUC, * $p$=0.0459; paired $t$ test. n=4 brain slices of 3 mice. Data shown as means ± SEM.

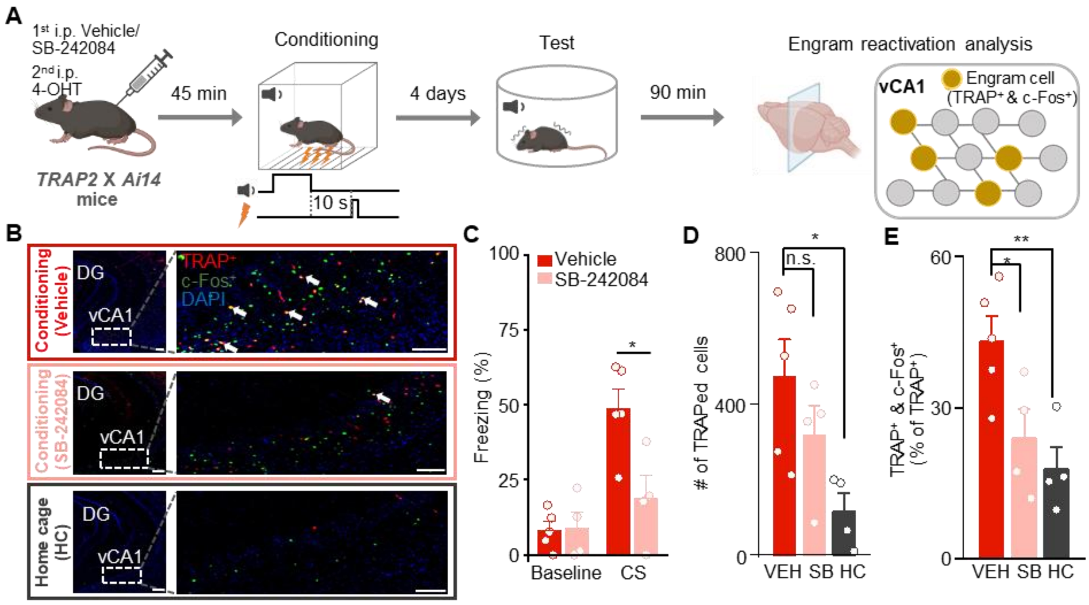


**Fig. S9. Identification of vCA1 engram cells during trace fear conditioning.**

(**A**) Experimental timeline: *TRAP2 x Ai14* mice received 10 mg/kg SB-242084 or VEH before 10-s interval fear conditioning. The mice were sacrificed within 90 min post-test. (**B**) Example images of TRAPed and c-Fos$^{+}$ neurons in the vCA1 following conditioning, conditioning with SB-242084 treatment, and in their home cage. White arrows indicate TRAP/c-Fos double-positive engram cells. (**C**) Summary of freezing behavior measured in the VEH and SB-242084 groups. * $p$=0.0217; unpaired $t$ test; n=4-5 mice/group. (**D**) Summary of the number of TRAPed cells in the indicated groups. * $p$=0.0213; One-way ANOVA with Dunnett's post hoc test; n=4-5 mice/group. (**E**) Summary of the percentage of TRAP/c-Fos double-positive cells among all TRAPed cells. ** $p$=0.0093, * $p$=0.0394; One-way ANOVA with Dunnett's post hoc test. Data shown as means ± SEM.
